# MEF2A is a negative regulator of β-Cell maturation and function

**DOI:** 10.64898/2026.03.09.710500

**Authors:** Ying Wang, Christine Darko, Tara Doma Lama, Andrew Rappa, Jeffery S. Tessem, Rohit B. Sharma

**Author notes:** Corresponding Author and Lead Contact Rohit B. Sharma, Assistant Professor, Director, Mouse Phenotyping Center, Weill Cornell Medical College, 413 E. 69^th^ Street, Belfer Research Building, BB616, New York, NY-10065; Correspondence.

## Abstract

Pancreatic beta cells produce and secrete insulin to maintain glucose homeostasis. Due to their high secretory activity, beta cells rely heavily on endoplasmic reticulum (ER) function and are particularly susceptible to ER stress, which contributes to beta cell dysfunction in diabetes. However, the transcriptional mechanisms linking ER stress to beta cell failure remain poorly understood. In this study, we investigated the role of the transcription factor Mef2a in ER stress-mediated beta cell dysfunction using primary mouse islet cells. ER stress induced by thapsigargin increased Mef2a expression and activated canonical unfolded protein response (UPR) pathways. Overexpression of Mef2a reduced beta cell proliferation, suppressed expression of key beta cell transcription factors including Pdx1, MafA, NeuroD1, and Nkx6.1, and impaired glucose-stimulated insulin secretion. Mef2a overexpression also altered mitochondrial respiration, characterized by reduced glucose-coupled respiration and increased maximal respiratory capacity. In contrast, Mef2a knockdown attenuated ER stress induced activation of ATF6 and IRE1/XBP1 dependent UPR genes. Importantly, reducing Mef2a expression preserved beta cell identity gene expression and improved insulin secretion during ER stress induced by thapsigargin or tunicamycin. Together, these findings identify Mef2a as a stress-responsive regulator that contributes to ER stress–mediated beta cell dysfunction and suggest that modulating Mef2a activity may help preserve beta cell function during metabolic stress.

## Introduction

Diabetes mellitus is characterized by dysfunction or loss of insulin-producing pancreatic β-cells [1]. In type 1 diabetes, this results from autoimmune destruction, whereas in type 2 diabetes (T2DM). β-cell failure occurs due to an inability to compensate for systemic insulin resistance [2; 3]. T2DM accounts for more than 90% of diabetes cases and affects over 500 million people worldwide [4]. Prediabetes represents an important window for intervention and is marked by insulin resistance, mild hyperglycemia, and compensatory Mechanisms[3; 4; 5; 6]. Increased insulin demand forces β-cells to elevate proinsulin synthesis, creating metabolic stress that can ultimately lead to β-cell dysfunction and loss [7; 8].

ER stress occurs when the ER’s protein-folding capacity is overwhelmed, leading to the accumulation of misfolded proteins and activation of the unfolded protein response (UPR). The UPR is mediated by three principal signaling branches: PERK, ATF6, and IRE1/XBP1, which collectively act to restore ER homeostasis by reducing protein synthesis, increasing expression of ER chaperones, and enhancing protein degradation pathways [9; 10]. In pancreatic β-cells, transient activation of the UPR can be adaptive, supporting increased insulin production during periods of metabolic demand. However, persistent ER stress leads to sustained UPR activation, which can impair β-cell identity, disrupt insulin secretion, and ultimately promote β-cell failure [11; 12; 13]. Both adaptive and overactive ER stress activate the same unfolded protein response (UPR) pathways [14]. Despite increasing recognition of the importance of ER stress in β-cell biology, the transcriptional regulators that link ER stress signaling to β-cell dysfunction remain poorly understood. Understanding the mechanisms that maintain ER homeostasis under metabolic stress may reveal new strategies to preserve β-cell function and prevent progression to diabetes.

Recent reports have shown that *Mef2a* is induced by chronic ER and metabolic stress in β-cells [15; 16; 17] and may play a role in β-oxidation, which impacts reactive oxygen species (ROS) and, in turn, causes ER stress. The MEF2 family of transcription factors (TFs) is vital for the proliferation, maturation, function, and survival of many cell types [18; 19; 20; 21]. *Mef2* has four family members, *Mef2a-d [21]*. Among the four family members, Mef2a and Mef2c are nearly identical, likely due to gene duplication. Mef2d is also similar to *Mef2a/c*, but it is the most evolutionarily divergent [18; 22]. Our unpublished RNA-seq data, supported by published datasets, show that *Mef2a* is the most abundant isoform in mouse and human β-cells, followed by *Mef2d* and *Mef2c* (expressed in mouse but absent in human). *Mef2b* is not expressed in the mouse and is least abundant in human β-cells [23; 24]. Ca2+-dependent kinases regulate MEF2 activity, CaMK [25], p38 MAPK [26], and calreticulin [27], which respond to intracellular Ca^2+^ fluctuations during ER stress but could be activated by several stress signaling inputs. ER stress disrupts Ca^2+^ homeostasis, depleting ER Ca^2+^ stores and altering cytosolic Ca^2+^ signaling [28; 29].

In this study, we investigated the role of Mef2a in ER stress–mediated β-cell dysfunction using primary mouse islet cells. We found that ER stress induces Mef2a expression and that increased Mef2a disrupts β-cell identity, mitochondrial metabolism, and insulin secretion. Conversely, reducing Mef2a expression attenuates UPR activation and preserves β-cell gene expression and insulin secretion during ER stress. Together, these findings identify Mef2a as a previously unrecognized regulator of ER stress signaling and β-cell functional integrity, providing new insight into transcriptional mechanisms that contribute to β-cell dysfunction during metabolic stress.

## Materials and Methods

### Mice

All mouse procedures were approved by the Weill Cornell’s Institutional Animal Care and Use Committee. C57BL6N mice were used for these studies. All mice were housed in a temperature-controlled room with a 12-h light/dark cycle and continuous access to standard mouse chow and water. Both male and female mice were used in all experiments at ages 15-60 weeks.

### Mouse islet isolation and cell culture

Islets were isolated by injecting collagenase into the common bile duct, followed by separating endocrine and exocrine tissue by Ficoll-Histopaque density gradient as described previously[30]. After resting overnight, islets were dispersed to single cells using 0.05% trypsin, plated on plastic for RNA and protein or on glass coverslips for immunostaining and microscopy, and cultured in islet complete media (ICM) containing RPMI 1640 with no glutamine and no glucose, 10% FBS, penicillin/streptomycin and supplemented with 15mM glucose in 24-well plates in a 37^°^C humidified incubator with 5% CO_2_. Dispersed mouse primary islet cells were treated with glucose or adenoviruses for 72 hr and then challenged with Thapsigargin and Tunicamycin for the last 24 hr of the culture. Adenovirus expressing *LacZ* (control), *Mef2a* (Mef2a overexpression), and Mef2a ShRNA (Mef2a knockdown) were used at a multiplicity of infection (MOI) of 5 for overexpression and 20 for knockdown of *Mef2a* [31]. Islet cells were harvested for molecular studies 72 hours after glucose and adenoviral treatment.

### Immunofluorescence

Islet cell cultures were fixed in 4% paraformaldehyde for 10 min at room temperature. For BrdU immunofluorescence, fixed cells or rehydrated paraffin sections were submerged in 1 N HCl for 25-30 min at 37°C, blocked for 2 hours in goat serum– based block with 0.1% Tween 20, labeled with primary antibodies, then secondary antibodies (Invitrogen), and mounted on glass slides with Fluoroshield mounting media containing DAPI (Sigma-Aldrich)[11; 32]. The primary antibodies used were guinea pig anti-insulin (catalog #A0564; Dako) and rat anti-BrdU (catalog # Ab6326; Abcam). Immunostained images were acquired using TissueGnostics NIKON microscope.

### Gene expression

For RNA, islet cells after culture were washed with ice-cold PBS, or liver, spleen, and muscle were suspended/homogenized in SKP buffer supplemented with 10% beta-mercaptoethanol from Norgen RNA/Protein isolation kit (Norgen, Canada). Total RNA was isolated from cells/tissues using the manufacturer’s protocol. 250-1000ng of total RNA underwent cDNA synthesis using the Superscript IV VILO kit from Invitrogen. Gene expression was measured by SYBR Green qPCR using primers previously used [31]. Data are expressed as ddCT (fold change) normalized to *Actb* and *Gapdh*.

### Immunofluorescence Image Quantification

Image data were quantified in an unbiased manner by blinding the images (by C.D., Y.W., and R.S.) Strataquest (TissueGnostics, USA). At least two images were manually checked post-quantification to confirm the data and assess for errors in the automated counts. 500-1000 β-cells were counted for each experimental condition. Mitochondrial Respiration using Seahorse XFe 96-well.

### Mitochondrial Respiration using Seahorse Assay

Mitochondrial respiration was measured in dispersed primary mouse islet cells using a Seahorse XF extracellular flux analyzer (Agilent) to determine the oxygen consumption rate (OCR). Cells were transduced with control adenovirus (*Ad-LacZ*) or *Ad-Mef2a* and plated in Seahorse XF microplates. On the day of the assay, cells were incubated in Seahorse XF assay medium (pH 7.4) containing **3** mM glucose and equilibrated for 1 h in a non-CO_2_ incubator. Basal respiration was measured, followed by sequential injections of 20 mM glucose, oligomycin, FCCP, and rotenone/antimycin A every 30 minutes to determine glucose-stimulated respiration, ATP-linked respiration, maximal respiration, and non-mitochondrial respiration. Data were analyzed using Seahorse Wave software, normalized to total DNA content and are presented as mean ± SEM.

### Glucose-Stimulated Insulin Secretion

Glucose-stimulated insulin secretion (GSIS) was assessed in dispersed primary mouse islet cells using a static incubation assay. Following the indicated treatments, cells were preincubated in Krebs-Ringer bicarbonate buffer containing low glucose (3 mM) for 1 h at 37°C. Cells were then sequentially incubated in buffer containing 3 mM glucose or 20 mM glucose for 30 min. Supernatants were collected after each incubation to measure secreted insulin. Cells were then lysed to determine total insulin content. Insulin secretion was quantified by ELISA and expressed either as secreted insulin normalized to total insulin content or as fold change relative to basal glucose conditions.

## Statistical Analysis

All the data are presented as mean ± SEM. The Statistical analyses were performed using Prism 10. A 2-tailed *unpaired* Student’s T-test calculated statistical significance (p-values) for comparing two groups or a one-way ANOVA for multiple comparisons. *P* < 0.05 was considered significant. All the representative replicates are plotted as individual data points or mentioned in the figure legends.

### Institutional Cores

We used the Mouse Phenotyping Center to run Seahorse assays.

## Results

### Mef2a and Mef2d are selectively activated by ER stress

To study the impact of ER stress on Mef2 family members, we treated mouse primary islet cells with Thapsigargin (Tg, 1 uM), a known ER stress inducer that depletes ER Ca stores by inhibiting SERCA2A [33]. Quantitative RT-PCR analysis revealed Tg treatment increased canonical ER stress indicators Grp78 and Ddit3 compared with DMSO-treated controls (Fig. 1A). Consistent with our previous findings Tg markedly upregulated the ATF6 target genes *Hyou1, HerpUD1*, and *Pdia4* (Fig. 1B). In addition, Tg stimulated the IRE1/XBP1pathway, as indicated by increased expression of the sXBP1 target genes *Erdj4, Ssr3*, and *Sec24d* (Fig. 1C). Recently ER stress and metabolic stress is shown to induce *Mef2a* in pancreatic β-cells [15; 16; 17]. We next examined whether ER stress influences the expression of the MEF2 family of transcription factors. Tg treatment resulted in a significant increase in *Mef2a* expression (∼2.8 fold). At the same time, *Mef2c* levels were not significantly altered, and *Mef2d* (∼1.2 fold) showed a modest induction (Fig. 1D). We also confirmed that *Mef2b* was not detected in mouse primary islet cells. Together, these results confirm robust activation of the UPR in response to Tg and identify Mef2a as a stress-responsive transcription factor that is selectively upregulated during ER stress in β-cells.

**Figure 1.**
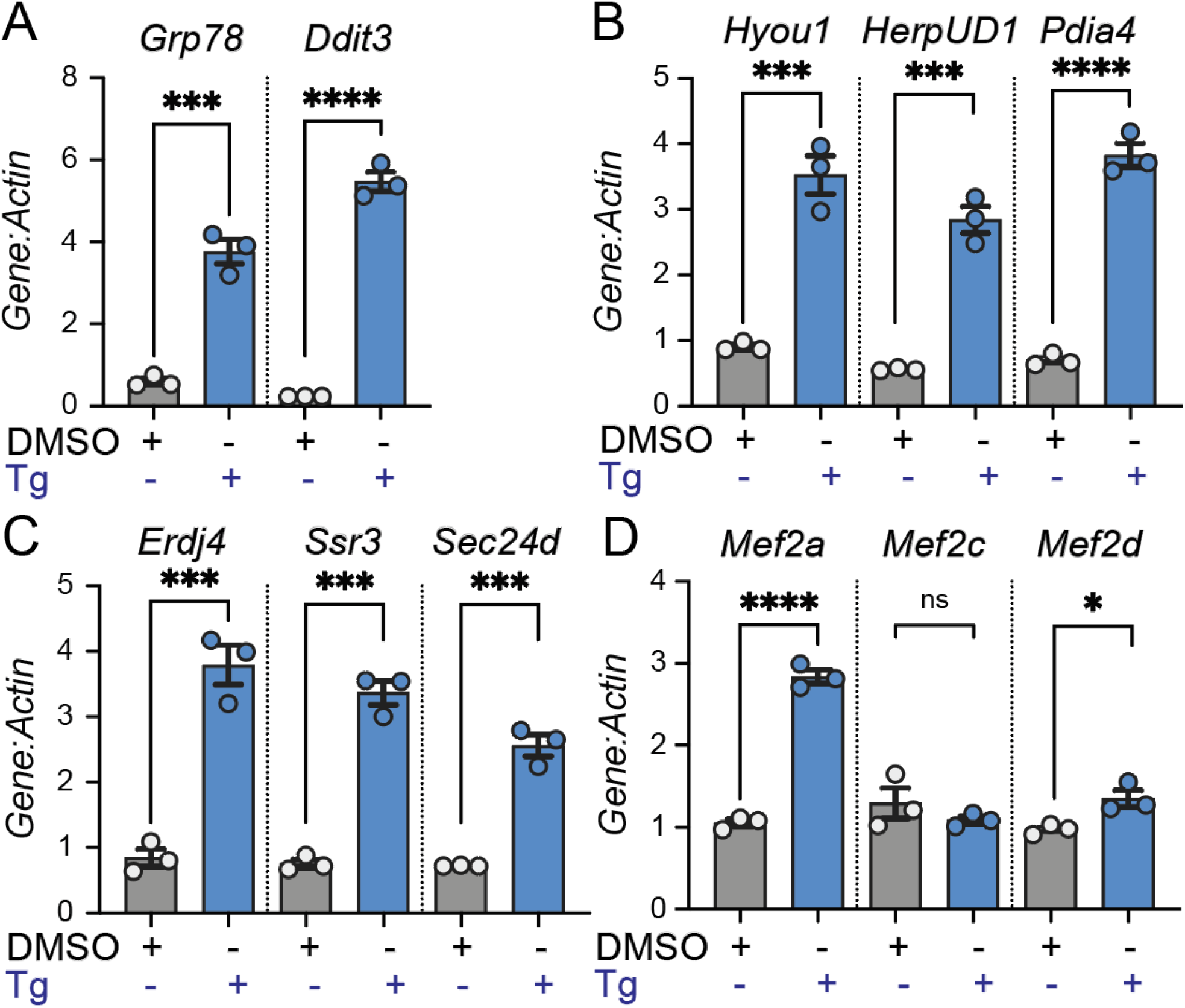
Thapsigargin induces ER stress response and activation of myocyte enhancer factor (Mef2) family. (A–D) RT–PCR analysis of gene expression in cells treated with vehicle (DMSO) or the ER stress inducer thapsigargin (Tg). (A) Classical ER stress markers *Grp78* and *Ddit3* are strongly induced by Tg treatment. (B–C) Tg also induces expression of ER stress response pathways, ATF6a measured by its transcriptional targets, *HerpUD1*, and *Pdia4*, and IRE1a pathway measured by sXBP1 transcriptional targets, *Erdj4, Ssr3*, and *Sec24d*. (D) Tg induces Mef2 family members, with *Mef2a* and *Mef2d* significantly, but *Mef2c* is not significantly changed. Data are presented as mean ± SEM with individual data points shown. Statistical significance was determined using an unpaired T-test (*p < 0.05, ***p < 0.001, ****p < 0.0001; ns, not significant).

### Mef2a overexpression suppresses β-cell proliferation, identity, and function

*Mef2* isoforms exhibit both functional specificity and redundancy. Mef2a and Mef2c are essential for myocyte differentiation and mitochondrial respiration[34]. Both Mef2a and Mef2d contribute to cardiomyocyte survival, with Mef2a compensating for the loss of Mef2d [35]. Mef2c is indispensable during embryonic development (whole-body Mef2c KO embryonic lethal); Mef2a can compensate for Mef2c function in adult tissues[19]. Collectively, these findings underscore the dominant role of Mef2a in myocytes, but its role in pancreatic β-cells has not been studied. To examine the functional role of MEF2A in pancreatic β-cells, dispersed islet cells were transduced with adenovirus expressing *Mef2a* (*Ad-Mef2a*) or control virus (Ad-LacZ), and the effects on β-cell identity gene expression, β-cell proliferation, and insulin secretion were assessed. Quantitative RT–PCR confirmed efficient overexpression of *Mef2a* in Ad-Mef2a–transduced cells compared with control cells under both low (5 mM) and high (15 mM) glucose conditions. While endogenous Mef2a expression in control cells was minimally affected by glucose, viral expression produced a robust increase in Mef2a mRNA, with the highest levels observed at 15 mM glucose (Fig. 2A) due to the glucose-responsive CMV promoter in the *Mef2a* overexpression virus. We next evaluated the effect of *Mef2a* overexpression on β-cell proliferation using BrdU incorporation assays. Immunofluorescence staining for insulin, BrdU, and DAPI revealed that high glucose (15 mM) significantly increased the number of proliferating β-cells in control virus– transduced islet cells compared with cells maintained at 5 mM glucose (Fig. 2B). In contrast, *Mef2a* overexpression markedly suppressed glucose-stimulated β-cell proliferation, as evidenced by a reduced number of insulin^+^/BrdU^+^ cells and a significant decrease in the percentage of BrdU-positive β-cells under high-glucose conditions (Fig. 2C). Because β-cell proliferation is often associated with changes in β-cell identity, we next examined the expression of key markers of β-cell maturation. Under high-glucose conditions (15 mM), Mef2a overexpression significantly reduced the expression of the β-cell identity genes *Pdx1, MafA, NeuroD1*, and *Nkx6*.*1*. In contrast, expression of *Ngn3* was not significantly altered (Fig. 2D). These findings indicate that elevated Mef2a levels disrupt the transcriptional network that maintains mature β-cell identity. Reduced expression of β-cell identity markers is associated with impaired β-cell function [36; 37; 38; 39; 40]. Finally, we assessed the functional consequences of *Mef2a* overexpression on glucose-stimulated insulin secretion (GSIS). In control cells, exposure to high glucose (20 mM) significantly increased insulin secretion compared with basal glucose (3 mM) (Fig. 2E). However, *Mef2a* overexpression exhibited a blunted secretory response to glucose, indicating impaired β-cell function. This defect was evident both in absolute insulin secretion and when secretion was normalized to total insulin content (Fig. 2E–F). Collectively, these results demonstrate that Mef2a overexpression negatively affects β-cell proliferation, reduces expression of key β-cell identity genes, and impairs glucose-stimulated insulin secretion, suggesting that elevated Mef2a levels compromise β-cell functional integrity.

**Figure 2.**
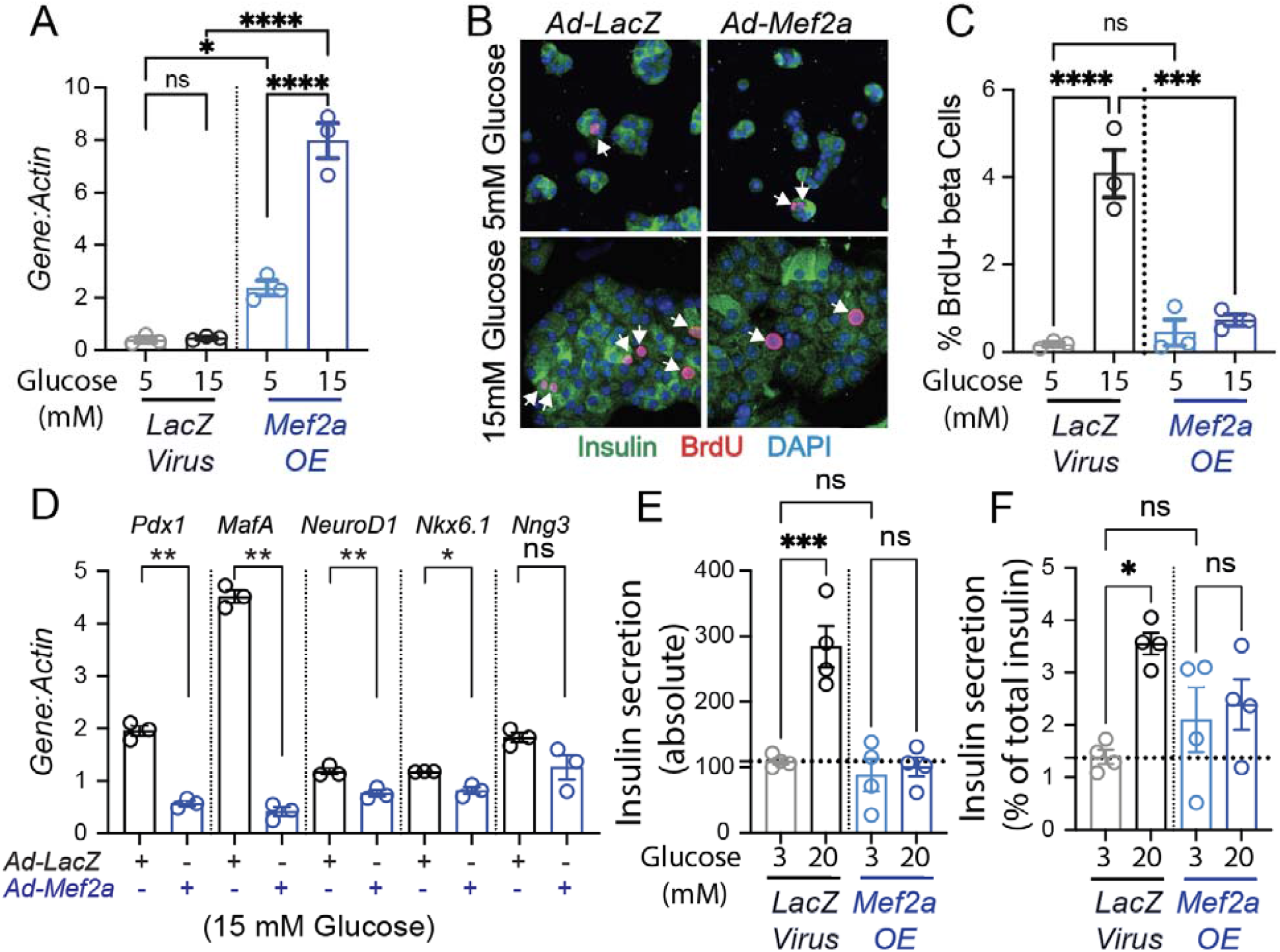
MEF2A overexpression impairs glucose-induced β-cell proliferation and alters β-cell maturation and insulin secretion. (A) RT–PCR analysis confirming Mef2a overexpression in mouse primary dispersed islet cells transduced with *Ad-Mef2a* compared with control *Ad-LacZ* virus under 5 or 15 mM glucose conditions. (B) Immunofluorescence images of islet cells cultured in 5 or 15 mM glucose and infected with *Ad-LacZ* or *Ad-Mef2a* and stained for insulin (green), BrdU (red), and DAPI (blue). Arrows indicate proliferating β-cells (insulin_+_/BrdU_+_, β-cells). (C) Quantification of the percentage of BrdU_+_ β-cells with *Mef2a* overexpression compared to the control condition. (D) RT–PCR analysis of β-cell identity and developmental transcription factors (*Pdx1, MafA, NeuroD1, Nkx6*.*1*, and *Nng3*) in islets infected with *Ad-LacZ* or *Ad-Mef2a* and cultured in 15 mM glucose. (E–F) Glucose-stimulated insulin secretion assays in islets infected with *Ad-LacZ* or *Ad-Mef2a* at 3 mM (low) and 20 mM (high) glucose. (E) Absolute insulin secretion. (F) Insulin secretion normalized to total insulin content. Data are presented as mean ± SEM with individual data points shown. Statistical significance was determined using unpaired *t*-tests for two-group comparisons or one-way ANOVA for multiple comparisons. Significance is indicated as *p* < 0.05, p < 0.01, *p < 0.001, **p < 0.0001; ns, not significant.

### Mef2a overexpression reduces basal and glucose-coupled mitochondrial respiration but enhances maximum respiration and spare capacity

As glucose-stimulated insulin secretion (GSIS) was impaired by *Mef2a*- overexpression, we next investigated whether mitochondrial metabolism was affected[41]. In pancreatic β-cells, mitochondrial oxidative metabolism plays a central role in coupling glucose sensing to insulin secretion. Following glucose uptake and glycolysis, pyruvate enters the mitochondria, where it fuels the tricarboxylic acid (TCA) cycle and oxidative phosphorylation, thereby generating ATP. The resulting increase in the ATP/ADP ratio closes ATP-sensitive K^+^ channels, triggering membrane depolarization, calcium influx, and insulin granule exocytosis[13; 42]. Consequently, defects in mitochondrial respiration are a well-established cause of impaired GSIS and β-cell dysfunction[43; 44]. Moreover, ER stress and transcriptional perturbations that disrupt β-cell identity have been shown to compromise mitochondrial metabolism and insulin secretory capacity[42; 45]. Given the reduced insulin secretion observed upon Mef2a overexpression, we hypothesized that mitochondrial oxidative metabolism might be impaired. Therefore, we assessed mitochondrial respiration by measuring oxygen consumption rate (OCR) to determine whether altered mitochondrial function contributes to the secretory defects observed after Mef2a overexpression. Basal respiration at 3 mM glucose was modestly reduced in Mef2a-overexpressing cells, although this difference was not significant (Fig. 3A–B). Upon stimulation with 20 mM glucose, control cells showed a clear increase in oxygen consumption rate (OCR), whereas Mef2a overexpression significantly reduced glucose-coupled respiration (Fig. 3A, C). Following FCCP treatment, maximal respiratory capacity and spare respiratory capacity were significantly increased in Mef2a-overexpressing cells (Fig. 3D–E). At the same time, ATP-linked respiration was not different (Fig. 3F). Non-mitochondrial respiration remained unchanged between groups (Fig. 3G). Together, these data indicate that Mef2a overexpression disrupts glucose-coupled mitochondrial respiration while increasing maximal respiratory capacity, suggesting altered mitochondrial metabolic regulation in β-cells, which could explain the decreased insulin secretion observed with Mef2a overexpression.

**Figure 3.**
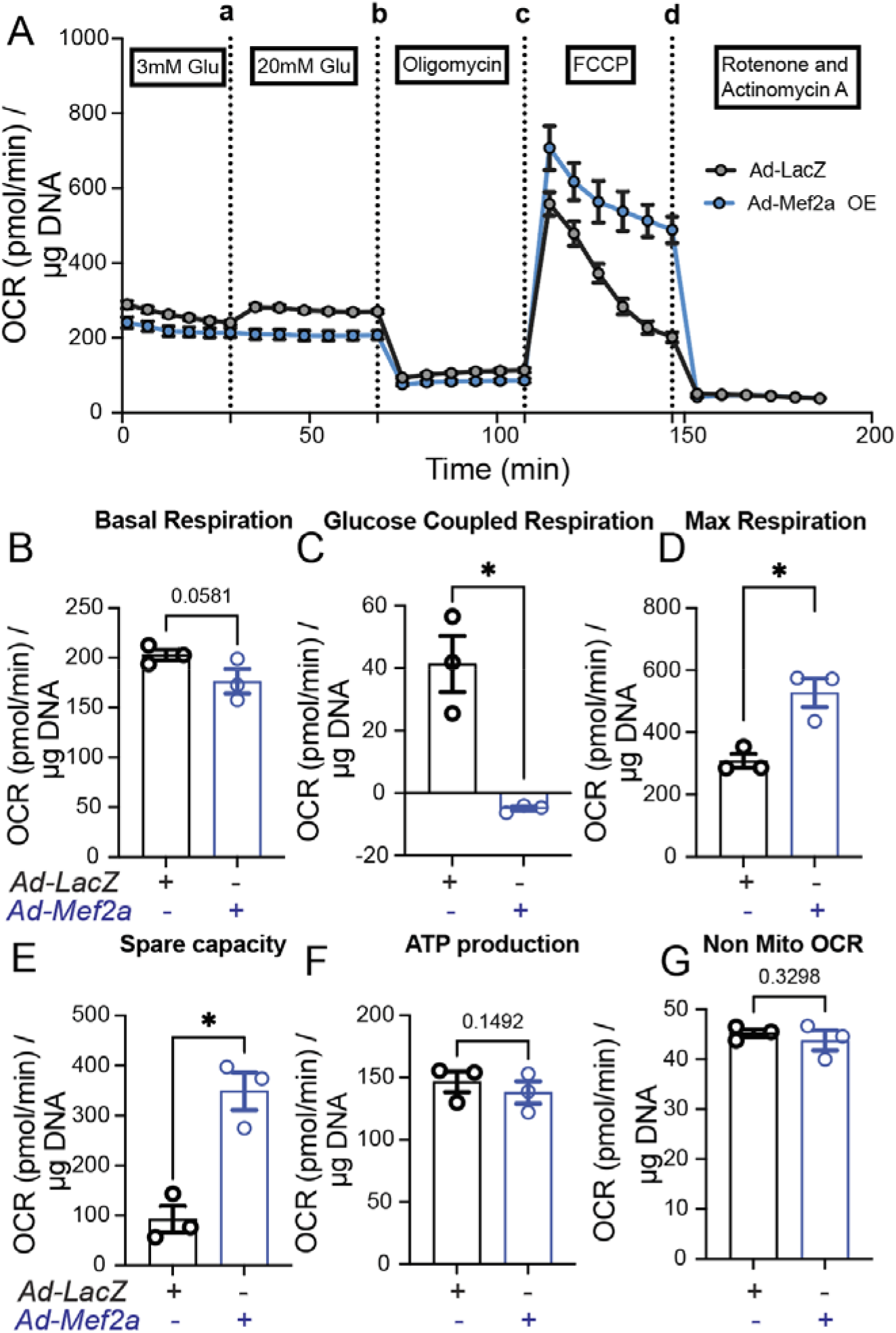
MEF2A overexpression reduces basal and glucose-coupled mitochondrial respiration but enhances maximum respiration and spare capacity in primary mouse islet cells. (A) Oxygen consumption rate (OCR) measured by Seahorse extracellular flux analysis in cells infected with control *Ad-LacZ* or *Ad-Mef2a* overexpression. OCR was monitored following sequential addition of 3 mM glucose, 20 mM glucose, oligomycin, FCCP, and rotenone/antimycin A, as indicated. Data are normalized to DNA content. (B–G) Quantification of mitochondrial respiration parameters derived from the Seahorse assay. (B) Basal respiration. (C) Glucose-coupled respiration. (D) Maximal respiration. (E) Spare respiratory capacity. (F) ATP production. (G) Non-mitochondrial OCR. Data are presented as mean ± SEM with individual data points shown. Statistical significance between two groups was calculated using an unpaired T-test and is indicated (*p < 0.05; ns, not significant).

### Mef2a knockdown attenuates UPR activation during ER stress

ER stress is a key contributor to β-cell dysfunction, particularly under conditions of increased insulin demand, where activation of the **unfolded protein response (UPR)** initially serves as an adaptive mechanism but can become maladaptive when chronically activated[9; 11; 14]. Mef2a expression increased during ER stress (Fig. 1), and Mef2a overexpression altered β-cell function (Figs. 2–3). We next examined whether Mef2a contributes to unfolded protein response (UPR) signaling. To determine whether Mef2a regulates UPR gene expression, dispersed islet cells were transduced with adenoviral shRNA targeting *Mef2a* (*Ad-Mef2a*) or control virus (*Ad-LacZ*). Under basal conditions, *Mef2a* knockdown had minimal effects on the expression of ER stress markers *Grp78* and *Ddit3*, as well as ATF6A target genes (*Hyou1, HerpUD1*, and *Pdia4*) and IRE1/XBP1 target genes (*Erdj4, Ssr3*, and *Sec24d*) (Fig. 4A–C), indicating that Mef2a is not required for basal UPR gene expression. We next examined gene expression following thapsigargin (Tg) induced ER stress, which robustly activates the UPR. Tg treatment markedly increased the expression of ATF6- and sXBP1-target genes in control cells. However, Mef2a knockdown significantly attenuated the induction of these UPR genes, including *Hyou1, HerpUD1, Pdia4, Erdj4, Ssr3*, and *Sec24d* (Fig. 4D–E). These findings suggest that Mef2a contributes to ER stress–induced activation of the UPR, and that reducing *Mef2a* levels dampens this transcriptional response. Because sustained UPR activation is known to promote β-cell dysfunction during chronic ER stress, the attenuation of ER stress signaling following *Mef2a* knockdown may represent a protective mechanism that helps preserve β-cell function under stress conditions.

**Figure 4.**
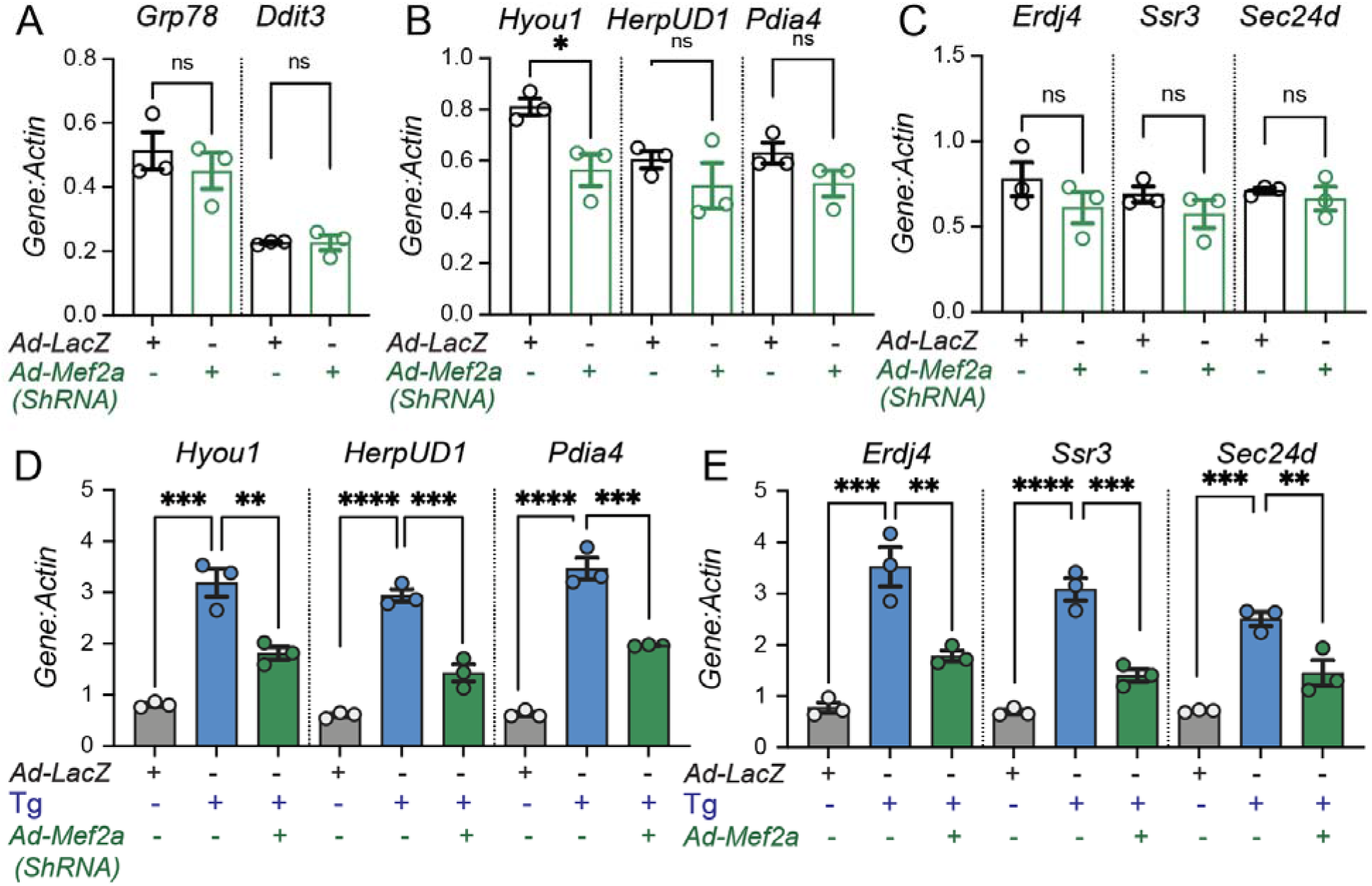
Mef2a knockdown has minimal impact on ER stress gene expression under basal conditions but attenuates UPR activation during thapsigargin-induced ER stress. (A–C) Quantitative RT–PCR analysis of ER stress–related genes following knockdown of *Mef2a* using adenoviral shRNA (*Ad-m-Mef2a*). Dispersed islet cells were transduced with control virus *Ad-LacZ* or *Ad-m-Mef2a*, and mRNA levels of general ER stress indicators Grp78 and Ddit3 (A), Atf6 transcriptional targets *Hyou1, HerpUD1*, and *Pdia4* (B), and IRE1 downstream sXbp1 transcriptional targets *Erdj4, Ssr3*, and *Sec24d* (C) were measured. Gene expression was normalized to *Actin*. Individual data points represent biological replicates. (D–E) Gene expression analysis following Thapsigargin (Tg) treatment with or without *Mef2a* knockdown. Cells were transduced with control virus (Ad-LacZ) or Ad-Mef2a in the presence of Tg. ATF6 target genes *Hyou1, HerpUD1*, and *Pdia4* (D) and sXBP1 target genes *Erdj4, Ssr3*, and *Sec24d* (E). Data are presented as mean ± SEM with individual data points shown. Statistical significance was determined using unpaired *t*-tests for two-group comparisons or one-way ANOVA for multiple comparisons. Significance is indicated as *p* < 0.05, p < 0.01, *p < 0.001, **p < 0.0001; ns, not significant.

### Mef2a knockdown preserves β-cell identity and insulin secretion under ER stress

To determine whether the reduced ER stress signaling observed with *Mef2a* knockdown protects β-cell function, we examined the expression of β-cell identity genes and insulin secretion following ER stress. Dispersed islet cells were transduced with control virus (*Ad-LacZ*) or *Ad-Mef2a shRNA* and treated with the ER stress inducers thapsigargin (Tg) or tunicamycin (Tm). Under Tg-induced ER stress, control cells exhibited a marked reduction in the expression of key β-cell transcription factors, including *Pdx1, MafA, NeuroD1*, and *Nkx6*.*1*, whereas *Mef2a* knockdown partially restored their expression (Fig. 5A). Similarly, treatment with Tm significantly suppressed β-cell identity genes in control cells, while Mef2a knockdown attenuated this effect, preserving the expression of *MafA, NeuroD1, Nkx6*.*1*, and Ngn3 (Fig. 5B). We next examined whether these transcriptional changes in β-cell genes were associated with improved β-cell function. Consistent with previous studies, ER stress significantly impaired glucose-stimulated insulin secretion (GSIS) in control islet cells treated with Tg or Tm[33; 46; 47]. Mef2a knockdown did not restore loss of glucose-stimulated insulin secretion in cells exposed to Tg (Fig. 5C); but significantly improved insulin secretion under ER stress caused by Tm (Fig. 5D). Together, these results demonstrate that reducing *Mef2a* expression mitigates ER stress–induced loss of β-cell identity and preserves insulin secretory function, supporting a role for *Mef2a* in promoting β-cell dysfunction during ER stress.

**Figure 5.**
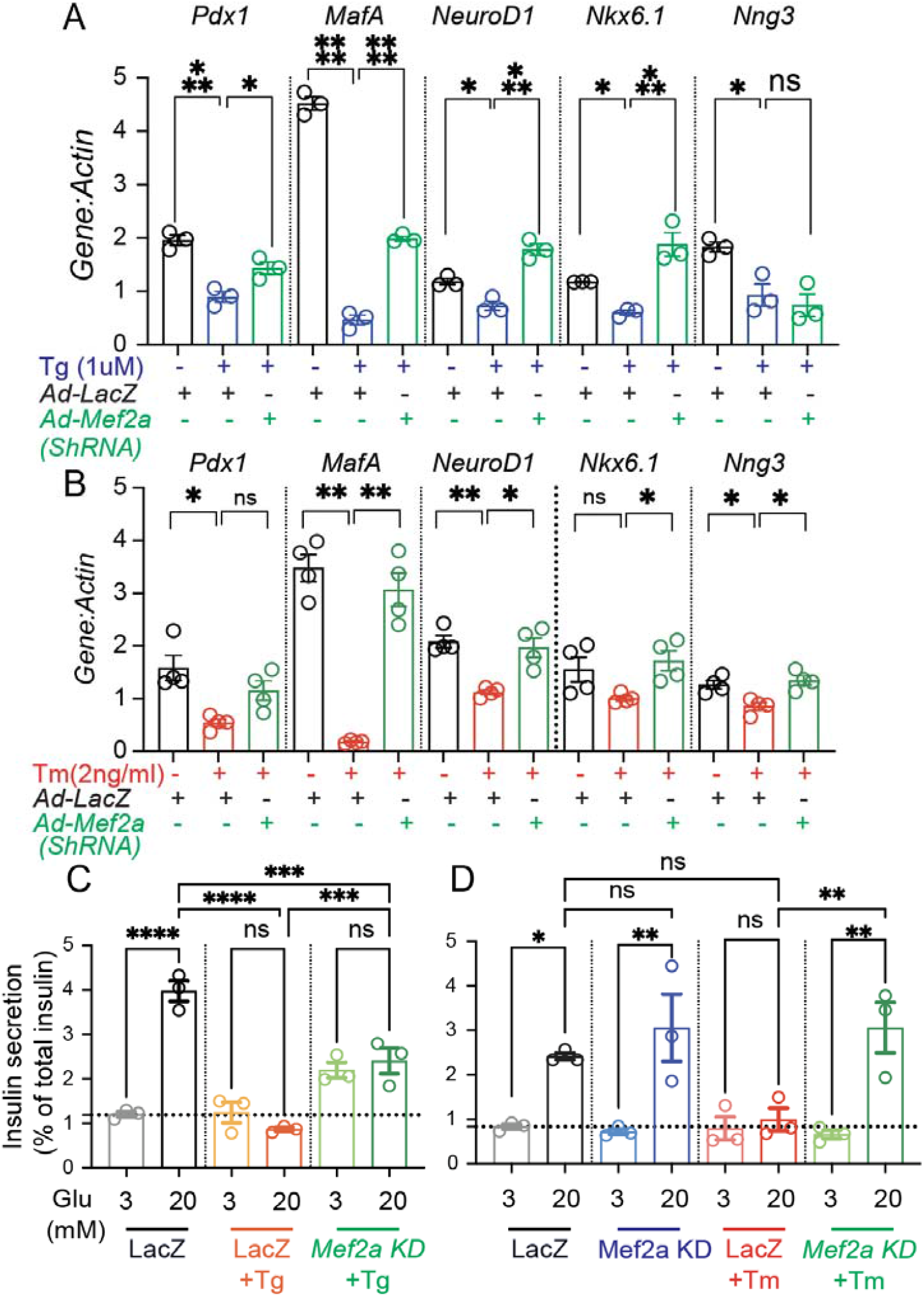
Mef2a knockdown preserves β-cell identity gene expression and insulin secretion during ER stress. (A) Quantitative RT–PCR analysis of β-cell transcription factors following thapsigargin (Tg, 1 μM) treatment with or without *Mef2a* knockdown. Dispersed islet cells were transduced with control adenovirus (Ad-LacZ) or adenoviral shRNA targeting *Mef2a* (*Ad-Mef2a*). mRNA levels of β-cell identity genes *Pdx1, MafA, NeuroD1, Nkx6*.*1*, and *Ngn3* were measured and normalized to Actin. (B) Gene expression analysis of the same β-cell transcription factors following tunicamycin (Tm, 2 ng/ml) treatment with or without *Mef2a* knockdown. (C–D) Glucose-stimulated insulin secretion (GSIS) assays were performed in dispersed islet cells transduced either with control virus (*LacZ*) or *Mef2a* shRNA. Cells were exposed to basal glucose (3 mM) or stimulatory glucose (20 mM). Insulin secretion was measured under control conditions or following Tg (C) or Tm (D) treatment. Secreted insulin is expressed as a percentage of total insulin content. Data are presented as mean ± SEM with individual biological replicates shown as dots. Statistical significance was determined using unpaired *t*-tests for two-group comparisons or one-way ANOVA for multiple comparisons. Significance is indicated as *p* < 0.05, p < 0.01, *p < 0.001, **p < 0.0001; ns, not significant.

## Discussion

Pancreatic β-cells are highly specialized secretory cells that rely on a robust endoplasmic reticulum (ER) and mitochondrial function to sustain insulin production and secretion. Consequently, β-cells are particularly vulnerable to ER stress, which contributes to β-cell dysfunction and loss in diabetes[48; 49]. In this study, we identify Mef2a as a stress-responsive transcription factor that promotes ER stress– induced β-cell dysfunction. Our findings demonstrate that ER stress induces Mef2a expression, that elevated *Mef2a* disrupts β-cell identity and mitochondrial metabolism, and reducing *Mef2a* dampens UPR activation and preserves β-cell function during ER stress.

The unfolded protein response functions to restore ER proteostasis by increasing chaperone expression, enhancing protein folding capacity, and reducing protein load[9; 10]. However, sustained or excessive activation of the UPR can become maladaptive and contribute to β-cell dysfunction[11; 12; 14]. The induction of Mef2a during ER stress suggests that this transcription factor may participate in transcriptional programs activated during cellular stress[16]. Functional experiments revealed that elevated *Mef2a* negatively affects β-cell proliferation, identity, function, and physiology.

Maintenance of β-cell identity depends on transcription factors such as PDX1, MAFA, NEUROD1, and NKX6.1, which regulate insulin gene expression and β-cell functional maturity, loss or reduction of these transcriptional regulators has been linked to β-cell dysfunction and dedifferentiation in diabetes [39; 40; 50; 51; 52; 53; 54]. Our findings suggest that increased Mef2a disrupts this transcriptional network and contributes to the loss of β-cell functional integrity.

In β-cells, glucose metabolism increases mitochondrial ATP production, which drives insulin secretion through closure of ATP-sensitive potassium channels and subsequent Ca^2+^ influx[41; 43]. We found that Mef2a overexpression impaired glucose-coupled respiration while increasing maximal and spare respiratory capacity. This observation suggests that the mitochondrial respiratory machinery remains intact but is functionally uncoupled from glucose metabolism, a defect associated with β-cell dysfunction in diabetes[42; 44]

To further understand the relationship between Mef2a and ER stress signaling, we examined the effects of *Mef2a* knockdown on UPR activation. Reducing Mef2a expression had minimal impact on basal ER stress gene expression but significantly attenuated the induction of ATF6- and IRE1/XBP1-target genes during thapsigargin-induced ER stress. These findings suggest that Mef2a functions as a transcriptional amplifier of ER stress signaling-induced β-cell dysfunction and dampening this response may help preserve β-cell function under stress conditions. Consistent with this concept, *Mef2a* knockdown partially preserved β-cell identity gene expression and improved insulin secretion during ER stress induced by thapsigargin but fully rescued these defects induced by tunicamycin. However, reducing Mef2a expression attenuated these effects, indicating that Mef2a contributes to ER stress– mediated β-cell dysfunction. These findings suggest that Mef2a acts downstream of ER stress to promote transcriptional and metabolic changes that compromise β-cell function.

A limitation of this study is that the experiments were performed in ex vivo dispersed mouse islet cells, which may not fully recapitulate the complex physiological environment of β-cells within intact islets or in vivo pancreatic tissue. β-cell interactions with other endocrine cell types, extracellular matrix components, and systemic metabolic signals may influence ER stress responses and β-cell function. Future studies using in vivo genetic models will therefore be important to determine the physiological role of Mef2a in β-cell adaptation and failure during metabolic stress.

Despite this limitation, a major strength of this study is the use of primary mouse islet cells rather than immortalized β-cell lines. Primary islets retain many of the physiological characteristics of native β-cells, including more accurate transcriptional programs, metabolic coupling mechanisms, and insulin secretion dynamics. In contrast, commonly used β-cell lines often exhibit altered metabolism and stress responses that may not fully represent native β-cell biology. Thus, studying Mef2a function in primary islet cells increases the physiological relevance of our findings.

In summary, our study identifies Mef2a as a novel regulator of ER stress signaling and β-cell dysfunction. We propose that ER stress induces *Mef2a* expression, thereby amplifying UPR transcriptional programs and disrupting mitochondrial metabolism and β-cell identity. Conversely, reducing Mef2a dampens ER stress signaling and preserves β-cell functional capacity. These findings provide new insight into how transcriptional regulators intersect with ER stress pathways to control β-cell health and suggest that modulating Mef2a activity may represent a potential strategy to protect β-cells during metabolic stress associated with diabetes.

## ACKNOWLEDGEMENTS AND FUNDING

The authors are thankful to members of the Joan and Sanford I. Weill Center for Metabolic Health for constructive feedback and discussion during work-in-progress meetings. This work is supported by startup funds from the Department of Medicine, Division of Endocrinology, Diabetes and Metabolism, Weill Cornell Medical College, NY to RBS.

## AUTHOR CONTRIBUTIONS

YW, CD, TDL, AR, and RBS performed the studies. JST and RBS conceptualized the study. RBS designed and conceptualized the experiments. RBS secured the funding. YW, CD, TDL, AR, and RBS performed the experiment. YW, CD, and RBS analyzed the data. RBS wrote the original draft, JST and RBS edited and finalized the manuscript. All authors had the opportunity to review the manuscript. RBS has full access to the data and takes full responsibility for its integrity.

## Bibliography

[1] G.C. Weir, and S. Bonner-Weir, Islet beta cell mass in diabetes and how it relates to function, birth, and death. Ann N Y Acad Sci 1281 (2013) 92–105.

[2] M. Cnop, N. Welsh, J.C. Jonas, A. Jorns, S. Lenzen, and D.L. Eizirik, Mechanisms of pancreatic beta-cell death in type 1 and type 2 diabetes: many differences, few similarities. Diabetes 54 Suppl 2 (2005) S97–107.

[3] E. Ferrannini, A. Natali, P. Bell, P. Cavallo-Perin, N. Lalic, and G. Mingrone, Insulin resistance and hypersecretion in obesity. European Group for the Study of Insulin Resistance (EGIR). J Clin Invest 100 (1997) 1166–73.

[4] A. American Diabetes, Diagnosis and classification of diabetes mellitus. Diabetes Care 37 Suppl 1 (2014) S81–90.

[5] E. Ferrannini, A. Gastaldelli, Y. Miyazaki, M. Matsuda, A. Mari, and R.A. DeFronzo, beta-Cell function in subjects spanning the range from normal glucose tolerance to overt diabetes: a new analysis. J Clin Endocrinol Metab 90 (2005) 493–500.

[6] A.G. Tabak, C. Herder, W. Rathmann, E.J. Brunner, and M. Kivimaki, Prediabetes: a high-risk state for diabetes development. Lancet 379 (2012) 2279–90.

[7] K.A. Goodge, and J.C. Hutton, Translational regulation of proinsulin biosynthesis and proinsulin conversion in the pancreatic beta-cell. Semin Cell Dev Biol 11 (2000) 235–42.

[8] J. Sun, J. Cui, Q. He, Z. Chen, P. Arvan, and M. Liu, Proinsulin misfolding and endoplasmic reticulum stress during the development and progression of diabetes. Mol Aspects Med 42 (2015) 105–18.

[9] P. Walter, and D. Ron, The unfolded protein response: from stress pathway to homeostatic regulation. Science 334 (2011) 1081–6.

[10] C. Hetz, K. Zhang, and R.J. Kaufman, Mechanisms, regulation and functions of the unfolded protein response. Nat Rev Mol Cell Biol 21 (2020) 421–438.

[11] R.B. Sharma, A.C. O’Donnell, R.E. Stamateris, B. Ha, K.M. McCloskey, P.R. Reynolds, P. Arvan, and L.C. Alonso, Insulin demand regulates beta cell number via the unfolded protein response. J Clin Invest 125 (2015) 3831–46.

[12] D.L. Eizirik, A.K. Cardozo, and M. Cnop, The role for endoplasmic reticulum stress in diabetes mellitus. Endocr Rev 29 (2008) 42–61.

[13] F.M. Ashcroft, and P. Rorsman, Diabetes mellitus and the beta cell: the last ten years. Cell 148 (2012) 1160–71.

[14] R.B. Sharma, H.V. Landa-Galvan, and L.C. Alonso, Living Dangerously: Protective and Harmful ER Stress Responses in Pancreatic beta-Cells. Diabetes 70 (2021) 2431–2443.

[15] T. Nammo, H. Udagawa, N. Funahashi, M. Kawaguchi, T. Uebanso, M. Hiramoto, W. Nishimura, and K. Yasuda, Genome-wide profiling of histone H3K27 acetylation featured fatty acid signalling in pancreatic beta cells in diet-induced obesity in mice. Diabetologia 61 (2018) 2608–2620.

[16] S. Khetan, S. Kales, R. Kursawe, A. Jillette, J.C. Ulirsch, S.K. Reilly, D. Ucar, R. Tewhey, and M.L. Stitzel, Functional characterization of T2D-associated SNP effects on baseline and ER stress-responsive beta cell transcriptional activation. Nat Commun 12 (2021) 5242.

[17] L.E. Parton, P.J. McMillen, Y. Shen, E. Docherty, E. Sharpe, F. Diraison, C.P. Briscoe, and G.A. Rutter, Limited role for SREBP-1c in defective glucose-induced insulin secretion from Zucker diabetic fatty rat islets: a functional and gene profiling analysis. Am J Physiol Endocrinol Metab 291 (2006) E982–94.

[18] M.J. Potthoff, and E.N. Olson, MEF2: a central regulator of diverse developmental programs. Development 134 (2007) 4131–40.

[19] N. Liu, B.R. Nelson, S. Bezprozvannaya, J.M. Shelton, J.A. Richardson, R. Bassel-Duby, and E.N. Olson, Requirement of MEF2A, C, and D for skeletal muscle regeneration. Proc Natl Acad Sci U S A 111 (2014) 4109–14.

[20] Y. Xiong, L. Wang, W. Jiang, L. Pang, W. Liu, A. Li, Y. Zhong, W. Ou, B. Liu, and S.M. Liu, MEF2A alters the proliferation, inflammation-related gene expression profiles and its silencing induces cellular senescence in human coronary endothelial cells. BMC Mol Biol 20 (2019) 8.

[21] B.L. Black, and E.N. Olson, Transcriptional control of muscle development by myocyte enhancer factor-2 (MEF2) proteins. Annu Rev Cell Dev Biol 14 (1998) 167–96.

[22] W. Wu, S. de Folter, X. Shen, W. Zhang, and S. Tao, Vertebrate paralogous MEF2 genes: origin, conservation, and evolution. PLoS One 6 (2011) e17334.

[23] M. Solimena, A.M. Schulte, L. Marselli, F. Ehehalt, D. Richter, M. Kleeberg, H. Mziaut, K.P. Knoch, J. Parnis, M. Bugliani, A. Siddiq, A. Jorns, F. Burdet, R. Liechti, M. Suleiman, D. Margerie, F. Syed, M. Distler, R. Grutzmann, E. Petretto, A. Moreno-Moral, C. Wegbrod, A. Sonmez, K. Pfriem, A. Friedrich, J. Meinel, C.B. Wollheim, G.B. Baretton, R. Scharfmann, E. Nogoceke, E. Bonifacio, D. Sturm, B. Meyer-Puttlitz, U. Boggi, H.D. Saeger, F. Filipponi, M. Lesche, P. Meda, A. Dahl, L. Wigger, I. Xenarios, M. Falchi, B. Thorens, J. Weitz, K. Bokvist, S. Lenzen, G.A. Rutter, P. Froguel, M. von Bulow, M. Ibberson, and P. Marchetti, Systems biology of the IMIDIA biobank from organ donors and pancreatectomised patients defines a novel transcriptomic signature of islets from individuals with type 2 diabetes. Diabetologia 61 (2018) 641–657.

[24] Q. Fu, H. Jiang, Y. Qian, H. Lv, H. Dai, Y. Zhou, Y. Chen, Y. He, R. Gao, S. Zheng, Y. Liang, S. Li, X. Xu, K. Xu, and T. Yang, Single-cell RNA sequencing combined with single-cell proteomics identifies the metabolic adaptation of islet cell subpopulations to high-fat diet in mice. Diabetologia 66 (2023) 724–740.

[25] H. Wu, F.J. Naya, T.A. McKinsey, B. Mercer, J.M. Shelton, E.R. Chin, A.R. Simard, R.N. Michel, R. Bassel-Duby, E.N. Olson, and R.S. Williams, MEF2 responds to multiple calcium-regulated signals in the control of skeletal muscle fiber type. EMBO J 19 (2000) 1963–73.

[26] M. Zhao, L. New, V.V. Kravchenko, Y. Kato, H. Gram, F. di Padova, E.N. Olson, R.J. Ulevitch, and J. Han, Regulation of the MEF2 family of transcription factors by p38. Mol Cell Biol 19 (1999) 21–30.

[27] A. Shalizi, B. Gaudilliere, Z. Yuan, J. Stegmuller, T. Shirogane, Q. Ge, Y. Tan, B. Schulman, J.W. Harper, and A. Bonni, A calcium-regulated MEF2 sumoylation switch controls postsynaptic differentiation. Science 311 (2006) 1012–7.

[28] S.R. Denmeade, C.M. Jakobsen, S. Janssen, S.R. Khan, E.S. Garrett, H. Lilja, S.B. Christensen, and J.T. Isaacs, Prostate-specific antigen-activated thapsigargin prodrug as targeted therapy for prostate cancer. J Natl Cancer Inst 95 (2003) 990–1000.

[29] J. Lytton, M. Westlin, and M.R. Hanley, Thapsigargin inhibits the sarcoplasmic or endoplasmic reticulum Ca-ATPase family of calcium pumps. J Biol Chem 266 (1991) 17067–71.

[30] J. Pascoe, D. Hollern, R. Stamateris, M. Abbasi, L.C. Romano, B. Zou, C.P. O’Donnell, A. Garcia-Ocana, and L.C. Alonso, Free fatty acids block glucose-induced beta-cell proliferation in mice by inducing cell cycle inhibitors p16 and p18. Diabetes 61 (2012) 632–41.

[31] R.B. Sharma, C. Darko, and L.C. Alonso, Intersection of the ATF6 and XBP1 ER stress pathways in mouse islet cells. J Biol Chem 295 (2020) 14164–14177.

[32] R.E. Stamateris, R.B. Sharma, Y. Kong, P. Ebrahimpour, D. Panday, P. Ranganath, B. Zou, H. Levitt, N.A. Parambil, C.P. O’Donnell, A. Garcia-Ocana, and L.C. Alonso, Glucose Induces Mouse beta-Cell Proliferation via IRS2, MTOR, and Cyclin D2 but Not the Insulin Receptor. Diabetes 65 (2016) 981–95.

[33] T. Aizawa, T. Yada, N. Asanuma, Y. Sato, F. Ishihara, N. Hamakawa, K. Yaekura, and K. Hashizume, Effects of thapsigargin, an intracellular CA2+ pump inhibitor, on insulin release by rat pancreatic B-cell. Life Sci 57 (1995) 1375–81.

[34] F.J. Naya, B.L. Black, H. Wu, R. Bassel-Duby, J.A. Richardson, J.A. Hill, and E.N. Olson, Mitochondrial deficiency and cardiac sudden death in mice lacking the MEF2A transcription factor. Nat Med 8 (2002) 1303–9.

[35] Y. Kim, D. Phan, E. van Rooij, D.Z. Wang, J. McAnally, X. Qi, J.A. Richardson, J.A. Hill, R. Bassel-Duby, and E.N. Olson, The MEF2D transcription factor mediates stress-dependent cardiac remodeling in mice. J Clin Invest 118 (2008) 124–32.

[36] D. Kawamori, H. Kaneto, Y. Nakatani, T.A. Matsuoka, M. Matsuhisa, M. Hori, and Y. Yamasaki, The forkhead transcription factor Foxo1 bridges the JNK pathway and the transcription factor PDX-1 through its intracellular translocation. J Biol Chem 281 (2006) 1091–8.

[37] W. Nishimura, T. Kondo, T. Salameh, I. El Khattabi, R. Dodge, S. Bonner-Weir, and A. Sharma, A switch from MafB to MafA expression accompanies differentiation to pancreatic beta-cells. Dev Biol 293 (2006) 526–39.

[38] J.C. Raum, K. Gerrish, I. Artner, E. Henderson, M. Guo, L. Sussel, J.C. Schisler, C.B. Newgard, and R. Stein, FoxA2, Nkx2.2, and PDX-1 regulate islet beta-cell-specific mafA expression through conserved sequences located between base pairs -8118 and -7750 upstream from the transcription start site. Mol Cell Biol 26 (2006) 5735–43.

[39] A.E. Schaffer, B.L. Taylor, J.R. Benthuysen, J. Liu, F. Thorel, W. Yuan, Y. Jiao, K.H. Kaestner, P.L. Herrera, M.A. Magnuson, C.L. May, and M. Sander, Nkx6.1 controls a gene regulatory network required for establishing and maintaining pancreatic Beta cell identity. PLoS Genet 9 (2013) e1003274.

[40] H. Wang, T. Brun, K. Kataoka, A.J. Sharma, and C.B. Wollheim, MAFA controls genes implicated in insulin biosynthesis and secretion. Diabetologia 50 (2007) 348–58.

[41] A. Wiederkehr, and C.B. Wollheim, Mitochondrial signals drive insulin secretion in the pancreatic beta-cell. Mol Cell Endocrinol 353 (2012) 128–37.

[42] E. Haythorne, M. Rohm, M. van de Bunt, M.F. Brereton, A.I. Tarasov, T.S. Blacker, G. Sachse, M. Silva Dos Santos, R. Terron Exposito, S. Davis, O. Baba, R. Fischer, M.R. Duchen, P. Rorsman, J.I. MacRae, and F.M. Ashcroft, Diabetes causes marked inhibition of mitochondrial metabolism in pancreatic beta-cells. Nat Commun 10 (2019) 2474.

[43] P. Maechler, and C.B. Wollheim, Mitochondrial function in normal and diabetic beta-cells. Nature 414 (2001) 807–12.

[44] D.G. Nicholls, The Pancreatic beta-Cell: A Bioenergetic Perspective. Physiol Rev 96 (2016) 1385–447.

[45] M. Szabat, M.M. Page, E. Panzhinskiy, S. Skovso, M. Mojibian, J. Fernandez-Tajes, J.E. Bruin, M.J. Bround, J.T. Lee, E.E. Xu, F. Taghizadeh, S. O’Dwyer, M. van de Bunt, K.M. Moon, S. Sinha, J. Han, Y. Fan, F.C. Lynn, M. Trucco, C.H. Borchers, L.J. Foster, C. Nislow, T.J. Kieffer, and J.D. Johnson, Reduced Insulin Production Relieves Endoplasmic Reticulum Stress and Induces beta Cell Proliferation. Cell Metab 23 (2016) 179–93.

[46] L. Zhao, H. Guo, H. Chen, R.B. Petersen, L. Zheng, A. Peng, and K. Huang, Effect of Liraglutide on endoplasmic reticulum stress in diabetes. Biochem Biophys Res Commun 441 (2013) 133–8.

[47] K. Tran, Y. Li, H. Duan, D. Arora, H.Y. Lim, and W. Wang, Identification of small molecules that protect pancreatic beta cells against endoplasmic reticulum stress-induced cell death. ACS Chem Biol 9 (2014) 2796–806.

[48] S.G. Fonseca, J. Gromada, and F. Urano, Endoplasmic reticulum stress and pancreatic beta-cell death. Trends Endocrinol Metab 22 (2011) 266–74.

[49] D.L. Eizirik, L. Pasquali, and M. Cnop, Pancreatic beta-cells in type 1 and type 2 diabetes mellitus: different pathways to failure. Nat Rev Endocrinol 16 (2020) 349–362.

[50] F.C. Pan, and C. Wright, Pancreas organogenesis: from bud to plexus to gland. Dev Dyn 240 (2011) 530–65.

[51] A. Swisa, B. Glaser, and Y. Dor, Metabolic Stress and Compromised Identity of Pancreatic Beta Cells. Front Genet 8 (2017) 21.

[52] S. Guo, C. Dai, M. Guo, B. Taylor, J.S. Harmon, M. Sander, R.P. Robertson, A.C. Powers, and R. Stein, Inactivation of specific beta cell transcription factors in type 2 diabetes. J Clin Invest 123 (2013) 3305–16.

[53] A.M. Vanhoose, S. Samaras, I. Artner, E. Henderson, Y. Hang, and R. Stein, MafA and MafB regulate Pdx1 transcription through the Area II control region in pancreatic beta cells. J Biol Chem 283 (2008) 22612–9.

[54] C.S. Hunter, M.A. Maestro, J.C. Raum, M. Guo, F.H. Thompson, 3rd, J. Ferrer, and R. Stein, Hnf1alpha (MODY3) regulates beta-cell-enriched MafA transcription factor expression. Mol Endocrinol 25 (2011) 339–47.

